# Dynein pulling forces on ruptured nuclei counteract lamin-mediated nuclear envelope repair mechanisms in vivo

**DOI:** 10.1101/138693

**Authors:** Lauren Penfield, Brian Wysolmerski, Reza Farhadifar, Michael Martinez, Ronald Biggs, Hai-Yin Wu, Michael Mauro, Curtis Broberg, Daniel Needleman, Shirin Bahmanyar

## Abstract

Recent work done exclusively in tissue culture cells revealed that the nuclear envelope (NE) undergoes ruptures leading to transient mixing of nuclear and cytoplasmic components. The duration of transient NE ruptures depends on lamins, however the underlying mechanisms and the relevance to *in vivo* events is not known. Here, we use *C. elegans* embryos to show that dynein forces that position nuclei increase the severity of lamin-induced NE ruptures *in vivo*. In the absence of dynein forces, lamin prevents nuclear-cytoplasmic mixing caused by NE ruptures. By monitoring the dynamics of NE rupture events, we demonstrate that lamin is required for a distinct phase in NE recovery that restricts nucleocytoplasmic mixing prior to the full restoration of NE rupture sites. We show that laser-induced puncture of the NE recapitulates phenotypes associated with NE recovery in wild type cells. Surprisingly, we find that embryonic lethality does not correlate with the incidence of NE rupture events suggesting that embryos survive transient losses of NE compartmentalization during early embryogenesis. In addition to presenting the first mechanistic analysis of transient NE ruptures *in vivo*, this work demonstrates that lamin controls the duration of NE ruptures by opposing dynein forces on ruptured nuclei to allow reestablishment of the NE permeability barrier and subsequent restoration of NE rupture sites.

## INTRODUCTION

The nuclear envelope (NE) is a large polarized double membrane sheet that serves to protect the genome in eukaryotic cells. The NE is continuous with the endoplasmic reticulum (ER) but has a unique protein composition and structure. In mitosis, the NE and associated proteins dramatically reorganize to promote NE breakdown (NEBD) and spindle microtubule capture of chromosomes. Following chromosome segregation, nuclear membranes emerge from the ER to reform around decondensing chromosomes (Hetzer 2010; Ungricht & Kutay 2017). A number of recent studies show that NE membrane dynamics are not restricted to NEBD and reformation (De Vos *et al.*, 2011; Vargas *et al.*, 2012; Denais *et al.*, 2016; Hatch and Hetzer, 2016; Webster *et al*., 2016). In migrating cells in culture, the NE undergoes reversible ruptures, which result in transient mixing of nuclear and cytoplasmic components, to promote passage of the nucleus through tight constrictions (Raab et al. 2016; Denais et al., 2016). In budding yeast, which undergo a closed mitosis, transient NE holes form at sites of defective nuclear pore complexes (NPCs) (Webster et al. 2016), the selective gatekeepers between the nucleus and cytoplasm (Hetzer & Wente 2009). The ESCRT-III machinery seals NE holes. ESCRT-III complex components are recruited to sites of defective NPCs (Webster et al. 2014; Webster et al. 2016), to NE ruptures in interphase (Denais *et al.*, 2016; Raab *et al.*, 2016), and to the reforming NE at the end of mitosis (Vietri et al. 2015; Olmos et al. 2015); further, inhibition of ESCRT-III complex components delays NE re-compartmentalization (Olmos et al. 2015; Vietri et al. 2015; Raab et al. 2016; Denais et al. 2016). Although NE ruptures are repaired and transient, they can adversely affect genomic integrity (Raab et al. 2016; Denais et al. 2016). This highlights the need to investigate mechanisms that induce NE ruptures, and promote recovery from ruptures, particularly in an *in vivo* system that can reveal consequences of this process in a physiologically relevant context.

Nuclear envelope ruptures occur at NE sites that are disrupted or deficient in nuclear lamins (Denais *et al*., 2016; Hatch and Hetzer, 2016; Raab *et al*., 2016; Robijns *et al*., 2016). The nuclear lamina is composed of metazoan-specific intermediate filament proteins lamin A/C and lamin B1/B2 and associated integral membrane proteins that include the conserved Lap2-Emerin-Man1 (LEM) family (Dechat et al. 2010; Wilson & Foisner 2010; Dechat et al. 2008). Lamins mechanically stabilize the NE by forming separate interdigitating meshworks that associate with the inner nuclear membrane facing chromatin (Davidson & Lammerding 2014; Shimi et al. 2015; Xie et al. 2016; Turgay et al. 2017; Dechat et al. 2008). In addition to mechanically supporting the nucleus, lamins and lamin-binding proteins regulate gene expression and organize chromatin (Reddy et al. 2008; Zullo et al. 2012; Mattout et al. 2015; Wilson & Foisner 2010), mediate DNA replication (Meier et al. 1991; Spann et al. 1997; Shimi et al. 2008; Moir et al. 2000), and impact spindle morphology (Tsai 2006; Zheng 2010; Ma et al. 2009; Goodman et al. 2010). Genetic disruption of lamin A/C and lamin-associated proteins leads to human disorders called laminopathies, which mostly affect tissues under high mechanical strain (Burke & Stewart 2013; Burke & Stewart 2006). Disease mutations in lamin A/C lead to nuclear deformations and loss of nuclear integrity in cells from diseased tissues (Burke & Stewart 2006; Burke & Stewart 2013). Recently, NE ruptures were reported in cultured fibroblasts from patients with lamin A/C mutations suggesting that this process may be relevant to lamin-associated diseases (De vos et al. 2011). An important next step is to understand if NE ruptures give rise to more severe NE integrity defects in diseased tissues. However, the low frequency and transient nature of NE ruptures make it challenging to investigate these events *in vivo*.

The reduction of each lamin subtype increases the frequency of NE ruptures in cancer cells in culture (Vargas et al. 2012; Denais et al. 2016; Raab et al. 2016, Irianto et al. 2016), highlighting the importance of lamins in preventing NE ruptures. Several lines of evidence suggest a critical role for lamin B1 in this process: reduction of lamin B1 results in NE regions devoid of nuclear lamina that correspond to rupture sites (Denais et al., 2016; Hatch and Hetzer, 2016), depleting lamin B1 results in the same frequency of NE ruptures as depleting all three lamins (Vargas et al. 2012), and local lamin B1 disruption at the NE has been implicated in NE sites of viral entry (de Noronha et al. 2001; Cohen et al. 2006). While these data support a specific role for lamin B1 in preventing NE ruptures, the reported structural interdependencies of lamins make it difficult to dissect individual roles for lamins in NE ruptures (Shimi et al. 2008). Also, it is challenging to separate lamins’ functions in transcriptional regulation and chromatin organization from its direct structural roles in NE integrity. The generation of double and triple lamin knockouts in mouse embryonic stem cells provides an instrumental tool for understanding essential roles for individual lamin subtypes (Kim et al. 2011; Kim et al. 2013), however the need to mechanistically dissect roles for lamins in NE rupture and repair mechanisms *in vivo* remains.

Our understanding of mechanisms that induce NE ruptures comes from recent studies in adherent cancer cells in culture that showed that, in addition to a role for lamins, actin-derived or mechanical compression forces imposed on the NE induce NE ruptures (Broers *et al*., 2004; Le Berre, Aubertin and Piel, 2012; Hatch and Hetzer, 2016). Inhibition of actin assembly rescues NE ruptures caused by lamin B1 depletion as detected by reduced cytoplasmic accumulation of GFP fused to a nuclear localization signal, an exclusively nuclear localized protein that accumulates in the cytoplasm upon NE rupture (Hatch and Hetzer, 2016). Actin inhibition also rescues the incidence of chromatin herniations, which form at rupture sites and correlate with rupture severity (Hatch and Hetzer, 2016). Consistent with a role for compression forces in inducing NE ruptures, constriction size and myosin activity influence the severity of migration-induced NE ruptures (Denais et al. 2016; Raab et al. 2016). Actin inhibition in lamin B1 depleted cells does not rescue the presence of NE regions devoid of lamin A, a hallmark of NE rupture sites, leading to the interpretation that disruption of lamins alone is not sufficient to induce ruptures (Hatch and Hetzer, 2016). Thus, the current model suggests that weakness in the nuclear lamina combined with cytoskeletal-generated nuclear pressure induces NE ruptures (Lammerding & Wolf 2016).

Repair of NE ruptures also depends on lamins (Vargas *et al*., 2012; Denais *et al*., 2016; Raab *et al*., 2016). The time between NE rupture and reestablishment of the nuclear permeability barrier depends on lamins (Vargas *et al*., 2012; Denais *et al*., 2016; Raab *et al*., 2016). NE rupture sites are devoid of lamins but lamin A/C accumulates to foci at rupture sites directly following migration-induced rupture and its levels of accumulation correlates with rupture severity, suggesting that it may play a role in repair of NE holes. However, how lamins mechanistically promote NE recovery following rupture is unknown.

Here, we show that the highly conserved single *Caenorhabditis elegans* lamin B isoform (Riemer et al. 1993; J Liu et al. 2000; Stuurman et al. 1998) restricts nucleocytoplasmic mixing following NE rupture by resisting dynein forces that position nuclei in the *C. elegans* zygote. Using an engineered hypomorphic allele in *C. elegans* lamin, which was designed based on a conserved disease-associated residue in lamin A/C (Fatkin *et al*., 1999), we reveal the dynamics of fluorescently-labeled NPCs, INM proteins and Histone2B at rupture sites during NE rupture and recovery. We show that the accumulation of the INM protein LEM-2, which acts as an adaptor for NE repair machinery, and loss of NPCs at NE rupture sites are independent of lamin and dynein; however, in the absence of dynein, lamin prevents nuclear entry of soluble GFP:α-tubulin upon NE rupture suggesting that lamin counteracts dynein induced tension to restrict nuclear-cytoplasmic mixing in response to NE rupture. The percentage of embryos that contain nuclei that undergo rupture does not correlate with the percentage of embryonic lethality indicating that NE rupture is tolerated during embryogenesis. Taken together, we discovered that lamin B promotes NE recovery from rupture by resisting dynein tension induced on ruptured nuclei and restricts loss of the nuclear permeability barrier in response to rupture, which we define as a distinct phase prior to full restoration of the NE. In addition, because the *C. elegans* zygote is transcriptionally quiescent (Edgar *et al*., 1994) these results represent direct structural functions for *C. elegans* lamin rather than secondary consequences of lamin inhibition on transcription.

## RESULTS

### A timeline of pronuclear positioning events in the *C. elegans* embryo to dissect structural roles for lamin

To investigate if *C. elegans* lamin protects against nuclear ruptures *in vivo*, we utilized the spatially and temporally defined pronuclear positioning events in the transcriptionally silent *C. elegans* zygote that contains a single lamin B gene (*lmn-1*). Prior to the first asymmetric division, dynein-dependent forces increase as the embryo progresses through the cell cycle to position the sperm- and oocyte-derived pronuclei (Cowan & Hyman 2004; Oegema & Hyman 2006). We monitored pronuclear position in embryos expressing GFP::Histone2B and GFP::γ-tubulin, to mark nuclei and centrosomes, respectively, to generate a timeline of pronuclear positioning events directly following oocyte fertilization through NE breakdown. The times of pronuclear positioning events are relative to pseudocleavage (PC) regression (0s, Figure 1A), a reference time point that coincides approximately with pronuclear meeting (~40s, Figure 1A). In our timeline, we incorporated information from other studies on the timing and location of active cytoskeletal forces and cell cycle transitions (Edgar and McGhee, 1988; O’Connell, 2000; Cowan and Hyman, 2004; Kimura and Onami, 2005; Oegema and Hyman, 2006; Portier *et al*., 2007; Dammermann et al., 2008).

**Figure 1.**
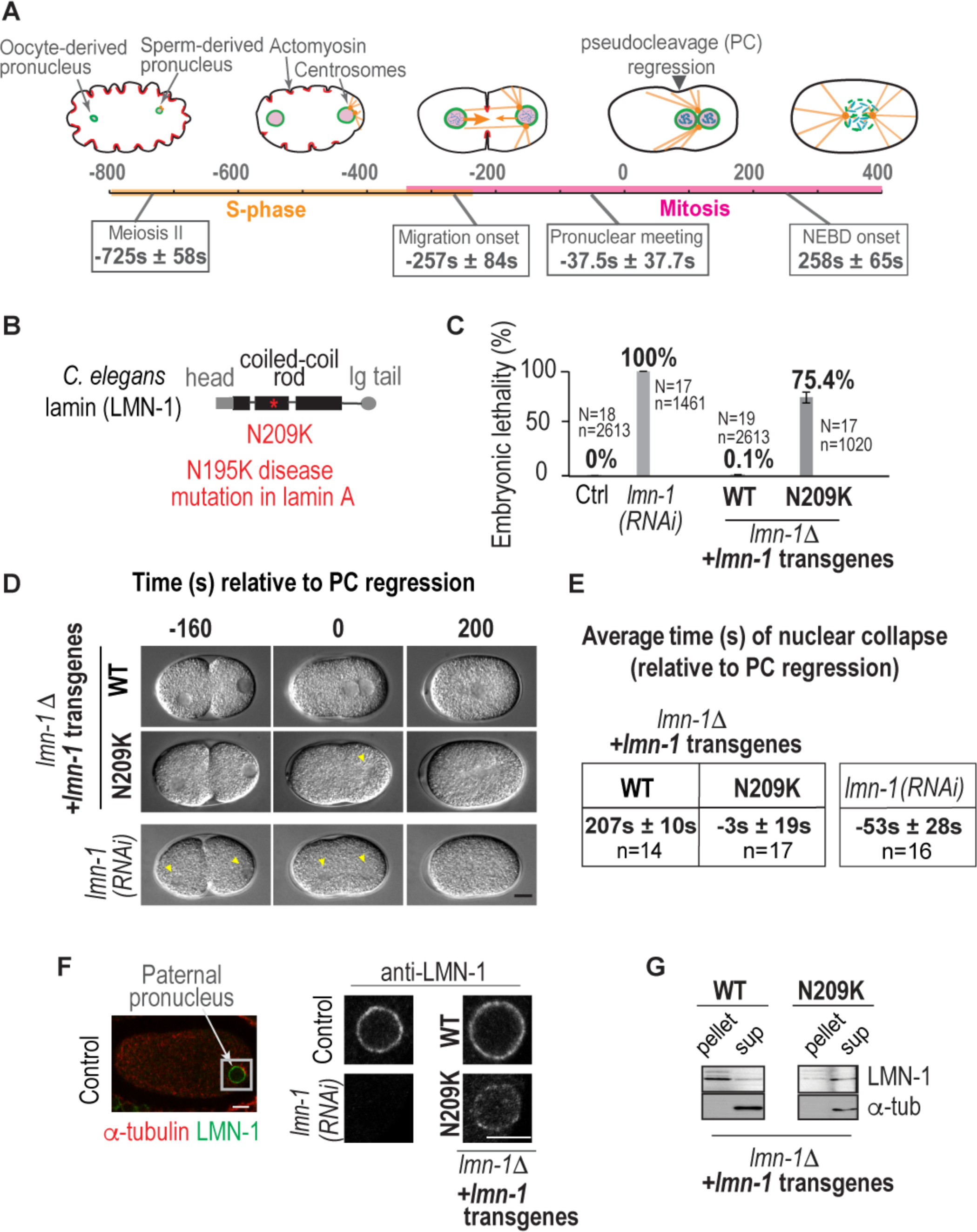
A disease-causing mutation in *C. elegans* lamin (lamin-N209K) is hypomorphic for lamin function. (A) Schematic illustrating the timeline of pronuclear positioning events in the one-cell stage *C. elegans* embryo relative to pseudocleavage (PC) regression. Times of key events are the average of 8 embryos ± S.D. (B) Schematic of *C. elegans* lamin protein domain structure and relative position of N209K disease mutation (red asterisk). (C) Percentage of embryonic lethality from the indicated conditions. N=number of worms, n=number of embryos. Data are represented as mean ± SEM. (D) DIC images from time-lapse series of *C. elegans* one-cell embryos for the indicated conditions. Arrowheads mark cleared regions that correspond to nuclear contents. Time points are in seconds relative to PC regression. Scale bar, 10μm. (E) Table showing the average times ± SEM of paternal nuclear collapse from DIC images in (D) for the indicated conditions. (F) Representative example of immunofluorescence image of a wild type one-cell embryo stained for lamin (red) and tubulin (green). Magnified image of paternal pronucleus immunostained with antibody against endogenous lamin from indicated conditions. Scale bars, 5μm. (G) Immunoblot probed for lamin (LMN-1) and α-tubulin after preparation of lysates into soluble (*sup*) and insoluble (*pellet*) fractions of worms from indicated genetic backgrounds. See also Figure S1.

Concurrent with anaphase of meiosis II of the oocyte (~−725s, Figure 1A), the paternal pronucleus and its attached centrosomes position adjacent to the plasma membrane at the opposite, future posterior end of the embryo (Cowan & Hyman 2004). At this time, chromosomes decondense and replicate as the embryo progresses through S-phase (Sonneville et al. 2015; Maddox et al. 2006). Centrosome maturation in S-phase leads to nucleation of short astral microtubules that facilitate centrosome separation to opposite sides of the paternal pronucleus via a dynein-dependent mechanism (~−450s to −250s; Gönczy *et al*., 1999; O’Connell, 2000; Malone *et al*., 2003; De Simone, Nédélec and Gönczy, 2016). Transition into prophase of mitosis (~−300s) induces chromosome condensation (Oegema & Hyman 2006). Elongated centrosome-nucleated microtubules reach the center of the embryo and associate with dynein motors anchored throughout the cytoplasm, which produces centering forces that pull on the paternal pronucleus (migration onset, ~−250s; Kimura and Onami, 2005). These forces position the paternal pronucleus away from the posterior edge (Kimura & Onami 2007; Kimura & Onami 2005). Microtubules reach NE associated dynein on the maternal pronucleus to facilitate pronuclear meeting (~−37s) (Cowan & Hyman 2004; Schmidt et al. 2005; Oegema & Hyman 2006; Malone et al. 2003). After pronuclear meeting (~−50s), centering, and rotation (~200s), NEBD proceeds in two phases (~250s): first, the dissociation of NPCs from nuclear membranes permeabilizes the NE allowing mixing of nuclear and cytoplasmic components, and, second, lamins disassemble as spindle microtubules penetrate nuclear membrane remnants to capture kinetochores attached to sister chromatids (Audhya et al. 2007; Portier et al. 2007). Coordination of NEBD and spindle assembly ensures mixing and congression of chromosomes from duplicated haploid genomes of maternal and paternal pronuclei prior to anaphase onset (~420s) (Bahmanyar et al. 2014; Lee et al. 2000; Oegema & Hyman 2006).

Thus, our detailed timeline of pronuclear positioning events in the *C. elegans* zygote offers the opportunity to investigate lamin’s function in maintaining nuclear integrity in the absence of transcription and in response to cytoskeletal forces that are relevant *in vivo*.

### A disease-causing mutation in a conserved residue in *C. elegans* lamin (lamin-N209K) is hypomorphic for lamin function

We reasoned that a hypomorphic allele in lamin would serve as a useful tool to investigate lamin’s structural roles in the early *C. elegans* embryo without potentially confounding effects of a penetrant lamin depletion. Using the MosI-mediated single copy insertion system (Frøkjær-Jensen et al. 2008), we engineered a *C. elegans* lamin transgene that carries a mutation in a conserved dilated cardiomyopathy linked residue in lamin A/C (N195K in human lamin A/C, N209K in *C. elegans* lamin; Figure 1B; (Fatkin *et al*., 1999; Wiesel *et al*., 2008). We chose this mutation because the same mutation in human lamin A reduces its incorporation into the nuclear lamina and decreases the ability of the NE to withstand mechanical forces (Zwerger et al. 2013). The untagged, RNAi-resistant wild type lamin transgene is expressed from its own *cis* regulatory elements (Figure S1A) and rescues the 100% embryonic lethality caused by depletion of endogenous lamin (Figure S1B, see Figure 1C for *lmn-1 (RNAi)*), thus validating this transgenic system as an effective method to compare wild type and mutant lamin alleles *in vivo*.

To more rigorously analyze the ability of engineered transgenes to support lamin’s functions, we used a lamin deletion strain (Figure S1C, referred to as *lmn-1Δ*), which is sterile but can be maintained as a heterozygous with a balancer (Figure S1D; Haithcock et al., 2005). The engineered lamin wildtype (WT-lamin) transgene supports embryo production and viability in the homozygous *lmn-1Δ* background (Figure 1C, S1D). The lamin-N209K mutant partially supports embryo production in the *lmn-1Δ* strain, and 75% of laid embryos do not hatch (Figure 1C, S1D). Because RNAi-depletion of lamin is 100% lethal (Figure 1C; Liu *et al*., 2000), the partial viability of embryos only expressing lamin-N209K upon RNAi-depletion of endogenous lamin (Figure S1B) or in the *lmn-1Δ* strain (Figure 1C) indicates that the lamin-N209K protein is partially functional.

Next, we performed time lapse imaging of the first embryonic division using differential interference contrast (DIC) microscopy to determine if the lamin-N209K mutant is able to support lamin’s functions in maintaining nuclear structure. WT-lamin pronuclei maintain their shape until NEBD (Figure 1D, 1E), whereas pronuclei in lamin RNAi-depleted embryos appear misshapen and barely visible prior to pseudocleavage regression (Figure 1D, 1E). The lamin-N209K transgene supports nuclear morphology until pseudocleavage regression at which time pronuclei lose their structure and become undetectable by DIC (Figure 1D, 1E). Thus, similar to its ability to partially support embryonic viability (Figure 1C), the lamin-N209K mutation partially supports lamin’s functions in maintenance of pronuclear integrity in the *lmn-1Δ* strain.

Prior work in *C. elegans* showed that a GFP-tagged lamin-N209K mutant protein expressed in the presence of endogenous lamin abnormally localizes to the NE (Wiesel et al. 2008). Our attempt to generate a functional fluorescent fusion WT-lamin transgene was unsuccessful (data not shown). Using an antibody we generated against *C. elegans* lamin, we localized WT and N209K-lamin transgenes in the *lmn-1Δ* background. Localization of lamin-N209K to the nuclear rim was significantly reduced in the absence of endogenous lamin (Figure 1F, S1E). Consistent with this, the lamin-N209K protein is more soluble under extraction conditions that do not normally solubilize lamin protein incorporated in the nuclear lamina (Figure 1G; Zwerger *et al*., 2013). Total lamin-N209K protein levels are reduced compared to the wild type transgene, however both transgenes express similar levels of lamin mRNA (Figure S1F, S1G). Endogenous lamin expressed from a single locus in the *het* deletion strain has reduced mRNA and protein levels, yet is normally localized to the nuclear rim and can support viable embryo production (“Het” in Figure S1D-G). Thus, the lamin-N209K mutant protein is intrinsically less able to incorporate into the nuclear lamina, which renders reduced localization to the nuclear rim and an unstable soluble pool of mutant protein.

Taken together, the functional and biochemical data support the conclusion that the lamin-N209K mutant protein is hypomorphic for lamin function, providing a useful tool to investigate more subtle structural functions for lamin during pronuclear positioning *in vivo*.

### Lamin and the lamin-N209 residue prevent transient and irreversible loss of the nuclear permeability barrier during distinct stages of pronuclear positioning

To dissect the underlying cause of nuclear phenotypes in lamin RNAi-depleted and lamin-N209K mutant embryos, we used time lapse imaging of embryos expressing fluorescently tagged markers. To generate strains that carry the lamin-N209K transgene and fluorescent markers, we took advantage of the engineered RNAi-resistant region in lamin transgenes (Figure S1A), which allows selective depletion of endogenous lamin. Because the lamin-N209K transgene supports 100% viability in the presence of endogenous lamin (data not shown), this strategy circumvents brood size and lethality issues that would otherwise hinder successful production of cross progeny.

Prior work showed that lamin RNAi-depletion inhibited the formation of an intact nuclear permeability barrier (Hayashi et al. 2012). To investigate the ability of the lamin-N209K mutant protein to support the NE permeability function for lamin, we monitored exclusion of soluble GFP::α-tubulin from the nuclear interior prior to pronuclear migration. GFP::α-tubulin was excluded from the paternal nuclear interior in embryos expressing the WT lamin transgene RNAi-depleted of endogenous lamin (hereafter referred to as WT-lamin) (Figure 2A-C). In contrast, in embryos expressing the lamin-N209K mutant protein RNAi-depleted of endogenous lamin (hereafter referred to as lamin-N209K) nuclear GFP::α-tubulin was present in 50% of paternal pronuclei prior to pseudocleavage regression (Figure 2A-C, Movie S1). 100% of embryos RNAi-depleted of lamin contained nuclear GFP::α-tubulin (Figure 2A-C), which is in agreement with prior studies (Hayashi, Kimura and Kimura, 2012) and our DIC imaging showing deformed pronuclei in these embryos at these time points (Figure 1D, 1E, S2). Quantification of the average nuclear GFP::α-tubulin fluorescence intensity during time points prior to pseudocleavage regression revealed that the maximum nuclear GFP::α-tubulin fluorescence intensity signal was similar between the different lamin-inhibition conditions indicating that GFP::α-tubulin rapidly equilibrates between the nucleus and cytoplasm upon entry (Figure 2A, 2B). The abrupt (within 20s) nuclear entry of GFP::α-tubulin was often followed by a gradual decrease GFP::α-tubulin in lamin-N209K and lamin RNAi-depleted pronuclei, and in some pronuclei repeated cycles of entry and partial exclusion of GFP::α-tubulin occurred (Figure 2A, 2B, Movie S1). To determine the percentage of nuclei that exclude GFP::α-tubulin following nuclear entry, we used 50% of the maximum nuclear signal as a threshold level of recovery from nuclear localization (also used by Vargas et al., 2012). Using this threshold, we found that 80% of lamin-N209K and 44% of lamin RNAi-depleted nuclei with nuclear GFP::α-tubulin exclude GFP::α-tubulin following its initial entry into the nucleus (Figure 2B, 2C, left plot). The time from entry to exclusion of nuclear GFP::α-tubulin lasted on average ~240s in lamin RNAi-depleted embryos and ~175s in lamin-N209K embryos (Figure 2B, 2C, Movie S1). Taken together, these data indicate that lamin-N209K and RNAi depletion of lamin impede the barrier function of the NE, allowing diffusion of soluble cytoplasmic GFP::α-tubulin into the nuclear interior; reduction of nuclear GFP::α-tubulin fluorescence following its entry indicates active export and thus recovery of NE barrier function. GFP::α-tubulin entered and persisted in the region of deformed and collapsed nuclei upon pronuclear migration and rotation in 100% of lamin RNAi-depleted and 60% of lamin-N209K embryos (Figure 2C, right plot; Figure S2). Thus, in contrast to transient breaches in the NE during early pronuclear positioning events, pronuclear meeting causes NEs to irreversibly collapse.

**Figure 2.**
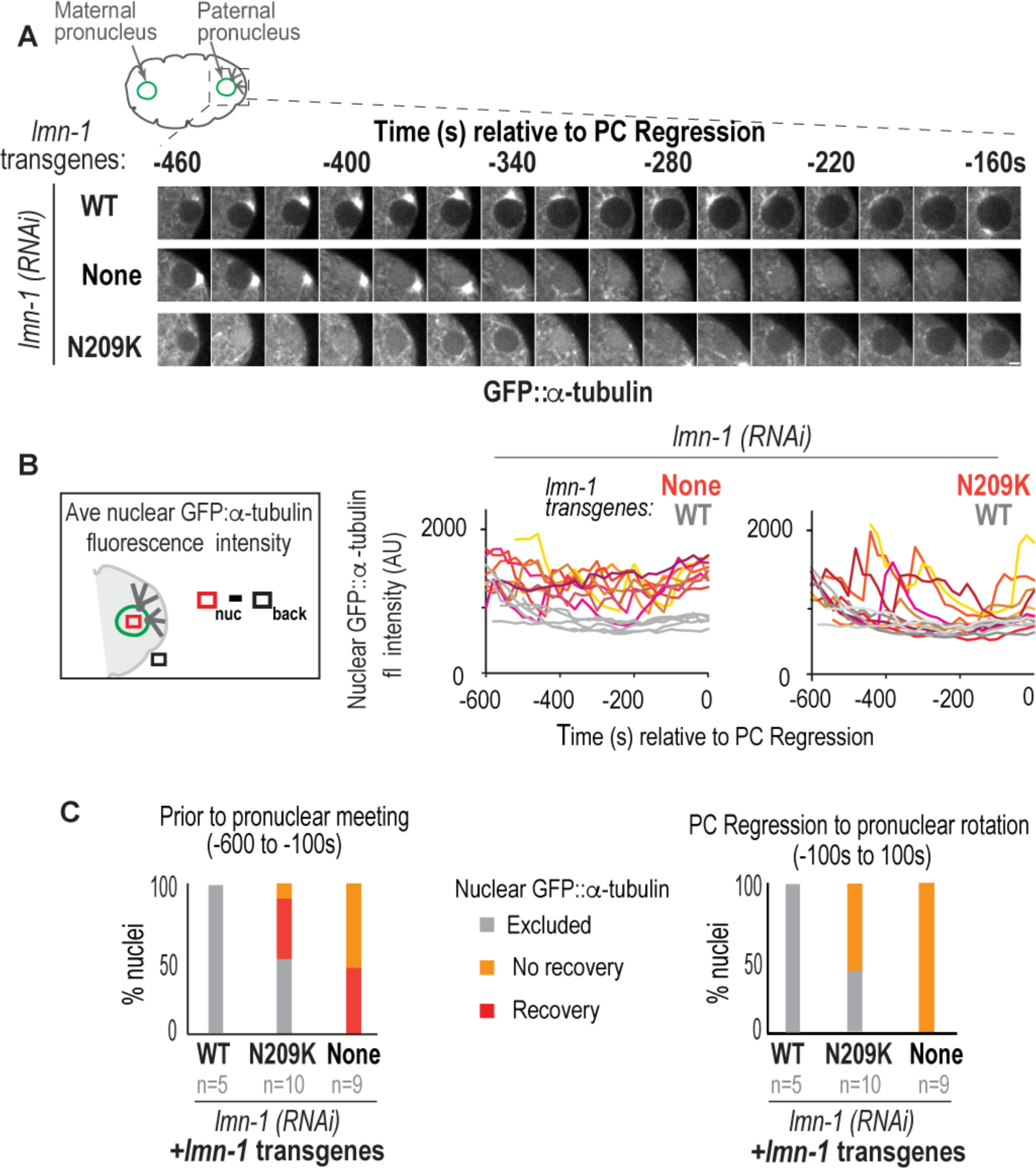
Lamin prevents transient nuclear entry of GFP::α-tubulin during pronuclear positioning. (A) Representative confocal images of a paternal pronucleus from time-lapse series of indicated conditions. Scale bar, 2.5μm. (B) (*Left*) Schematic of method used to measure nuclear GFP::α-tubulin fluorescence intensity levels. Average intensity of a 2.3 × 2.3μm box in the nucleus (red box) was subtracted by the average background intensity of an identical region placed in an area outside of embryo (black box, *back*). (*Right*) Plots of average nuclear GFP::α-tubulin fluorescence intensities at indicated time points relative to PC regression from indicated conditions. Each trace represents intensity measurements of one paternal pronucleus. Grey traces on both graphs correspond to wild type paternal pronuclei (C) Plot of percent of paternal nuclei in which GFP::α-tubulin is excluded (grey), nuclear GFP::α-tubulin that recovers to <50% of maximum level of nuclear fluorescence intensity (red), and nuclear GFP::α-tubulin that does not recover to <50% of maximum level of nuclear fluorescence intensity (orange). (*Left*) Plot represents data from measurements performed prior to pronuclear migration (−600 to −200s). Averages of time of recovery of GFP:α-tubulin exclusion: 240s ± 35s in lamin RNAi-depleted, 175s ± 33s for lamin-N209K, n=4 ruptured nuclei. (*Right*) Plot represents measurements performed at pronuclear migration until pronuclear rotation (−100s to 100s). n=number of embryos. Plots summarize data shown for each condition in Figure 2B. See also Figure S2 and Movie S1.

We conclude that lamin and the N209 residue in lamin prevent transient loss of the nuclear permeability barrier when dynein forces are relatively weak prior to pseduocloeavage regression. When dynein forces imposed on nuclei are relatively strong to facilitate pronuclear meeting lamin and the N209 residue in lamin protect against irreversible loss of the nuclear permeability barrier.

### Lamin and the lamin-N209K mutation induce nuclear envelope ruptures that result in nuclear envelope and chromatin reorganization at rupture sites

We predicted that transient loss of the nuclear permeability barrier in lamin-inhibited embryos is likely a consequence of NE ruptures. In mammalian cells, transient ruptures occur in regions that are devoid of nuclear pore complexes (NPCs) and lamins (De vos et al. 2011; Denais et al. 2016; Raab et al. 2016). To determine if pronuclei in lamin-inhibited embryos displayed NE regions devoid of NPCs, we imaged Nup160::GFP, a member of the NPC scaffold, and mCherry:Histone2B to mark chromatin (Hattersley et al. 2016). In S-phase, WT-lamin paternal pronuclei contain decondensed chromatin surrounded by a smooth continuous ring of Nup160::GFP (−420s in Figure 3A, 3B; Movie S2). In 69% of lamin-N209K and 100% of lamin RNAi-depleted embryos, a distinct gap in the Nup160::GFP signal at the nuclear rim is present at time points that correspond to S-phase and early prophase (Figure 3A-C; ~−600s to −380s in Figure 3D; Movie S2). The Nup160::GFP gap first appears during the time range of transient loss of the nuclear permeability barrier observed by nuclear GFP::α-tubulin (see Figure 2). Thus, lamin inhibition causes a gap in Nup160::GFP at the nuclear rim that likely marks the site of NE rupture.

**Figure 3.**
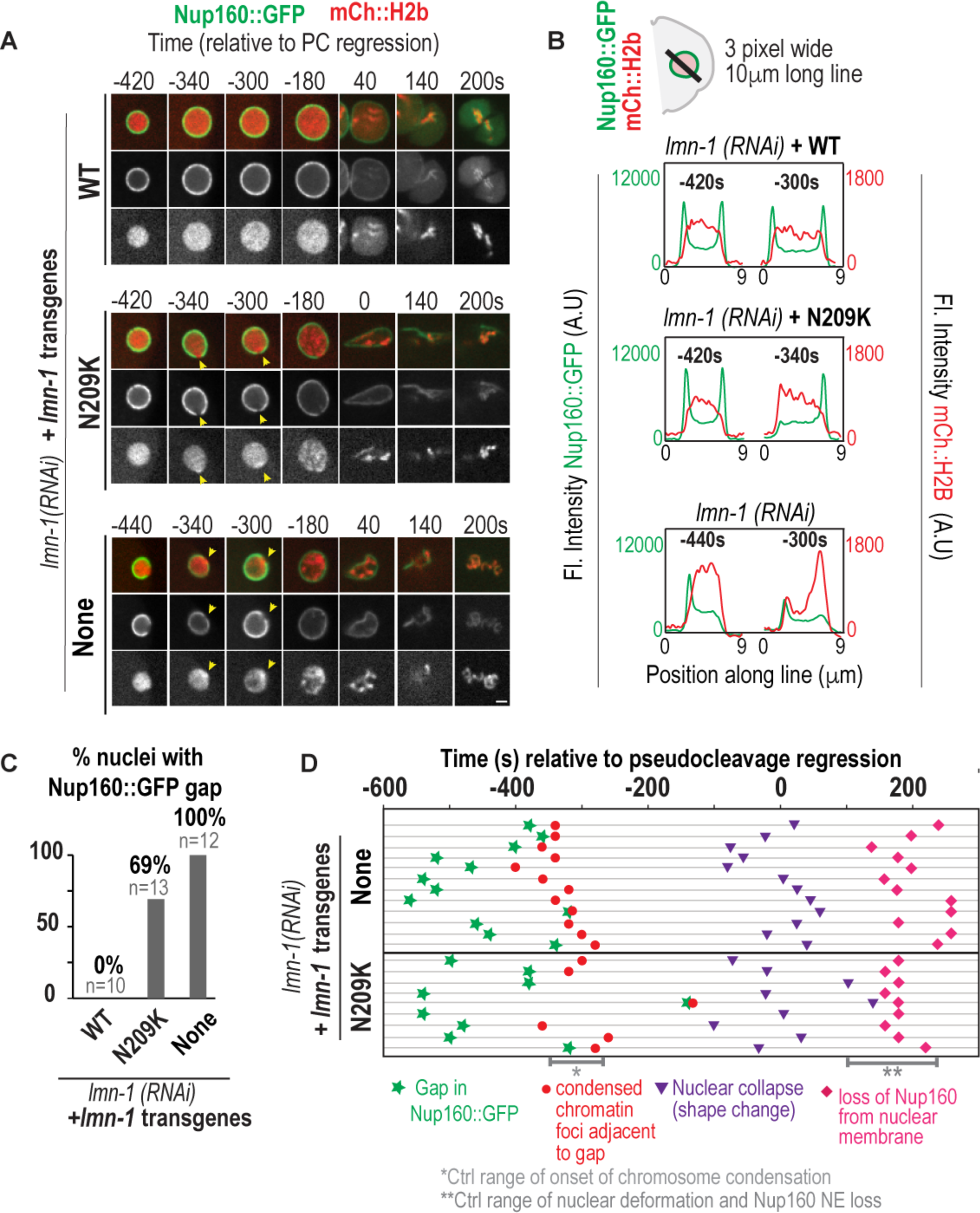
Nup160::GFP gaps precede mitotic-entry coupled chromatin reorganization at the nuclear envelope rupture sites. (A) Select representative confocal images of paternal pronuclei from time-lapse series of embryos expressing Nup160::GFP and mCherry::Histone2B from indicated conditions. Scale bar, 2.5μm. Times are in seconds and relative to PC Regression. Arrowheads in middle and bottoms panels marks site of gap in Nup160::GFP at nuclear rim and focus of condensed mCherry::Histone2B.. (B) Plots represent fluorescent intensities of Nup160::GFP (green) and mCherry::Histone2B (red) along a 3 pixel wide lines drawn across nucleus shown in (A) at indicated times relative to PC Regression. (C) Plot of percent of paternal pronuclei containing a Nup160::GFP gap from indicated conditions. n=number of pronuclei. (D) Timeline of events scored from confocal timelapse images of individual paternal pronuclei (horizontal lines) from indicated conditions. Time is in seconds relative to PC regression. Grey brackets with single asterisk indicates the range of time in which control pronuclei initiate chromosome condensation as measured in Figure 1A. Double asterisk represents the range of time in which control paternal pronuclei undergo nuclear deformation and loss of Nup160::GFP from the NE. See also Movie S2.

Subsequent to the appearance of a Nup160::GFP gap in the nuclear rim, and coincident with entry into mitotic prophase, a highly condensed spot of mCherry:Histone2B appears precisely in the region devoid of Nup160::GFP in lamin-N209K and lamin-RNAi depleted conditions (−340s and −300s in Figure 3A, 3B; ~−380s and −300s in Figure 3D; Movie S2). Following its initial appearance at this site, in lamin RNAi-depleted embryos chromosomes become highly compacted and localize directly adjacent to the nuclear periphery (−180s in Figure 3A). This phenotype is less severe in lamin-N209K embryos: discrete condensed chromosome puncta distribute throughout the nuclear interior at time points in which control pronuclei do not contain distinct chromosomes (−180s in Figure 3A, Movie S2). Thus, in both lamin-N209K expression and lamin RNAi-depletion, a condensed chromatin focus appears at NE rupture sites in early prophase, which is directly followed by abnormal nuclear distribution of prematurely compacted chromosomes.

As expected, upon pronuclear migration, lamin-N209K and lamin RNAi-depleted NEs marked by Nup160::GFP deform and collapse onto chromatin (Figure 3A, 0 to 140s; Figure 3D, −100 to 100s; Movie S2). Dissociation of Nup160::GFP from NE remnants in lamin-N209K and lamin RNAi-depleted nuclei occurs approximately 200s after pseudocleavage regression, which corresponds to the timing of cell-cycle regulated NEBD in control embryos (Figure 3A, 200s; Figure 3D, 200s; Movie S2). Nup160::GFP accumulates onto kinetochores, the attachment sites for spindle microtubules, with similar timing in control and lamin-inhibited conditions (Figure 3A, ~140s). Thus, defects in nuclear structure and chromatin condensation caused by lamin inhibition in early embryos are not a consequence of abnormal cell cycle progression.

Taken together, our time-lapse analysis show that initial ruptures in the NE occur in S-phase as marked by a clearance of Nup160::GFP at the nuclear rim. NE rupture induces condensed chromatin puncta at rupture sites upon entry into prophase, which is followed by global defects in chromatin condensation and organization.

### Nuclear envelope reorganization following lamin- or laser-induced NE rupture

We predicted that if NE regions devoid of Nup160::GFP are sites of NE ruptures, then they should be transient and resolve with similar kinetics as reestablishment of the NE permeability barrier, assayed by recovery of nuclear GFP:α-tubulin following its entry (Figure 2). At time points prior to pseudocleavage regression, 100% of Nup160::GFP gaps resolve in lamin-N209K nuclei and 83% resolve in lamin RNAi-depleted nuclei (Figure 4A, 4B, Movie S2). We found that the lifetimes of Nup160::GFP gaps, measured from their initial appearance to their resolution, were ~190s in lamin RNAi-depleted embryos and ~130s in lamin-N209K embryos (Figure 4C, S3A), similar to the duration of initial entry of nuclear GFP::α-tubulin to 50% recovery (~240s in lamin RNAi-depleted embryos and ~175s in lamin-N209K embryos, Figure 2B, C). Furthermore, concurrent with appearance of a Nup160::GFP gap in the nuclear rim, we observed defects in nuclear shape that recovered as Nup160::GFP gaps resolved (Figure 4A). Thus, these data indicate that, in agreement with prior work in cancer cells (De Vos *et al*., 2011; Denais *et al*., 2016; Raab *et al*., 2016), NPCs clear at sites of NE rupture. We show that Nup160::GFP gaps resolve with the same kinetics as reestablishment of the NE permeability barrier (Figure 2B, 4C, S3A) and correlate with recovery of nuclear shape defects (Figure 4A). However, we observe re-entry of GFP::.a-tubulin into the nucleus during or after 50% recovery as determined by a 20% increase from the previous time point during the recovery process (33% (3/9) for lamin RNAi-depleted and 40% (n=5) for lamin-N209K had reentry of GFP::α-tubulin) (Figure 2A-B), yet only a single Nup160::GFP gap in 72% of lamin RNAi-depleted and 92% of lamin-N209K nuclei suggesting that repeated breaches in the permeability barrier likely occur at the same NE rupture site. Furthermore, while the duration of entry of nuclear GFP:α-tubulin to 50% recovery is similar to the kinetics of NPC gap appearance and resolution, nuclear GFP:α-tubulin levels decrease directly following entry suggesting that the permeability barrier function of the NE is recovered despite lack of NPC reinsertion at NE rupture sites during that time frame. Together, our results are consistent with the roles of lamins in limiting the duration of NE recovery (Vargas *et al*., 2012; Denais *et al*., 2016; Raab *et al*., 2016), but here, we elucidate that lamin promotes recovery by restricting access to the nucleoplasm at NE rupture sites.

**Figure 4.**
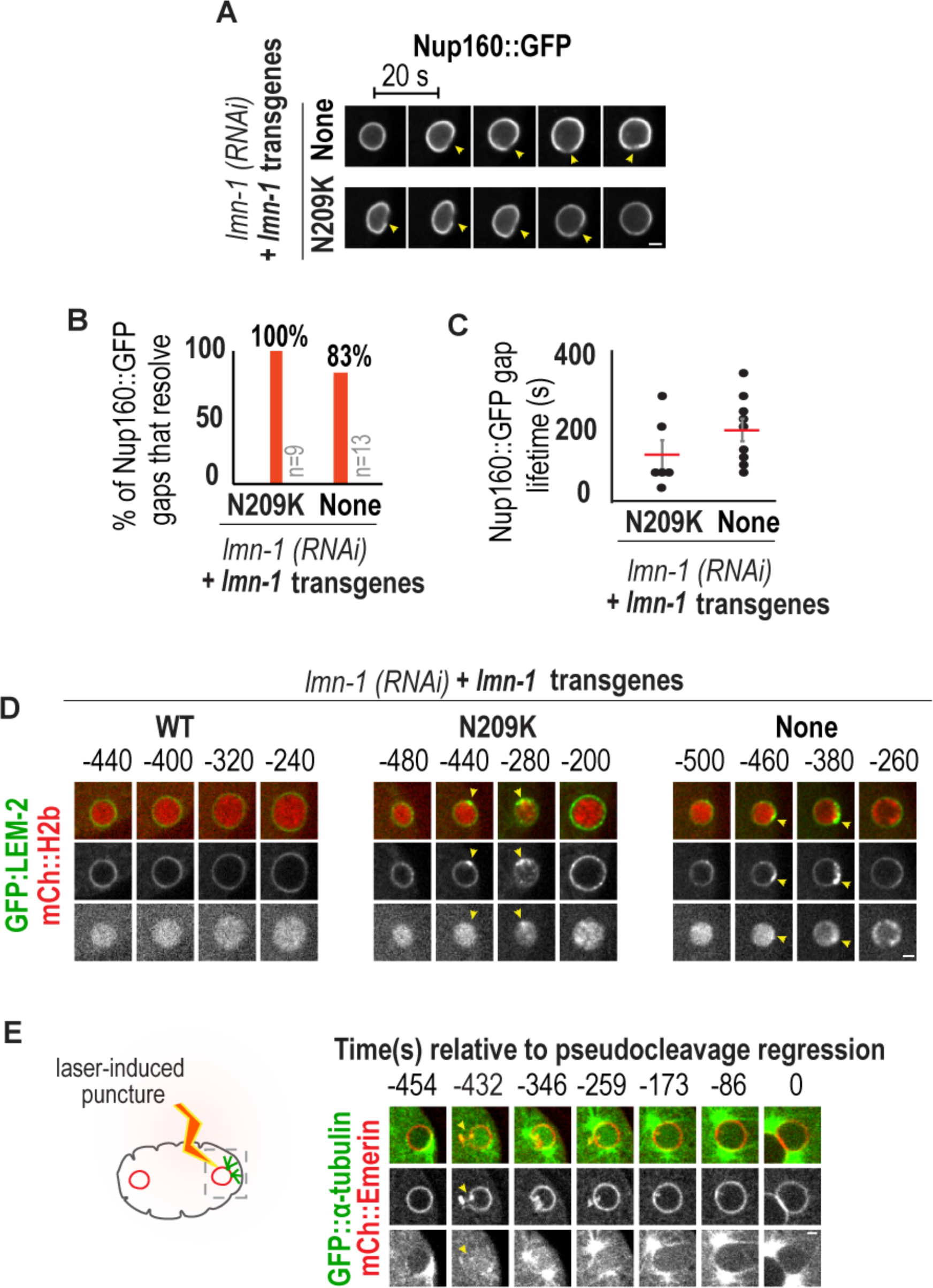
Nuclear envelope repair from ruptures is lamin-independent and occurs in wild type embryos following laser-induced puncture of the nuclear envelope. (A) Consecutive confocal images from 20s interval time series of Nup160::GFP expressing paternal pronuclei from indicated conditions. Arrowhead marks region of gap in Nup160::GFP signal at nuclear rim. Scale bar, 2.5μm. (B) Percent of Nup160::GFP gaps that resolve after appearance in the indicated conditions. n= number of embryos. (C) Plot of individual times (black) and the average time (red) ± SEM in seconds between appearance and resolution of Nup160::GFP gap. Average gap lifetimes: 191s ± 29s, lamin RNAi-depletion; 127s ± 38s, lamin-N209K. (D) Select confocal images of paternal pronuclei labeled with GFP::LEM-2 (green) and mCherry::histone (red). Arrowhead marks distinct region in nuclear rim with GFP::LEM-2 accumulation. Time is in seconds relative to PC regression. Scale bar, 2.5μm. (E) Select confocal images of time-lapse series following laser-induced puncture of mCherry::Emerin labeled NE (see −432s). Simultaneous imaging of GFP::α-tubulin to show the breach and recovery of the NE permeability barrier in wild type embryos. Time is in seconds relative to PC Regression. Scale bar, 2.5μm. See also Figure S3 and Movie S2 and Movie S3.

To further support the idea that NE regions devoid of Nup160::GFP that are adjacent to condensed chromatin puncta mark transient NE rupture sites, we imaged the inner nuclear membrane protein GFP::LEM-2. In mammalian cells, LEM2 acts as the nuclear adaptor that recruits ESCRT-III repair machinery to seal NE openings at the end of mitosis, and this function for LEM-2 is conserved in yeast (Gu et al. 2017). We found that GFP::LEM-2 accumulates in distinct regions in the nuclear rim ~500s prior to pseudocleavage regression in lamin-N209K and lamin RNAi-depleted pronuclei (Figure 4D), which corresponds to the time of nuclear entry of GFP::α-tubulin (Figure 2) and the appearance of a Nup160::GFP gap in the nuclear rim (Figure 3A, 3D). Further, condensed chromatin puncta appear directly adjacent to the site of GFP::LEM-2 accumulation at the nuclear rim (Figure 4D). Because LEM2 has a conserved function in recruiting repair machinery to sites of NE holes in other systems (Webster et al. 2016; Gu et al. 2017), we conclude that sites of LEM-2 accumulation and subsequent dispersal are NE rupture sites undergoing repair. In the majority of embryos (86% (n=7) of lamin-RNAi depleted and 75% (n=4) of lamin-N209K embryos), the LEM-2 punctum appeared at one site and persisted at this site over time (Figure 4D).

To recapitulate NE repair in wild type pronuclei, we monitored nuclear GFP::α-tubulin and NE dynamics after laser-induced puncture of the NE. We focused on time points that correspond to NE ruptures in lamin-inhibited embryos. Using an 800nm Titan Sapphire laser, we generated 4μm by 1μm by 0.5μm and 4μm by 2μm by 0.5μm punctures in the nuclear rim marked by mCherry::Emerin. We simultaneously monitored nuclear entry of GFP::α-tubulin to ensure a successful breach in the nuclear permeability barrier and to monitor NE recovery. Similar to transient lamin-induced NE ruptures, laser-induced puncture caused nuclear deformation and rapid equilibration of nuclear and cytoplasmic soluble GFP::α-tubulin followed by recovery of both nuclear deformation and entry of GFP::α-tubulin (Figure 4E, S3B, Movie S3). Laser punctured pronuclei recovered from nuclear shape changes, and 40% recovered to 60% of the maximum nuclear GFP::α-tubulin signal (Figure 4E, Figure S3C). The duration of nuclear localized GFP::α-tubulin was ~300s, which is ~125s and ~60s longer than the average duration of lamin-N209K and lamin-induced ruptures (Figure 2B), respectively, to recovery of 50% of the maximum nuclear GFP::α-tubulin signal. The range of recovery times between these different conditions suggests that the time to recovery correlates with the size of NE holes and that these are relatively smaller in lamin-N209K expressing embryos.

Overall, the laser-induced punctured nuclei recapitulate phenotypes associated with lamin-dependent nuclear rupture and repair in wild type embryos, and show that active NE repair mechanisms are in place even in nuclei that are not prone to NE ruptures or loss of integrity. We conclude that the duration of loss of the nuclear permeability barrier depends on the severity of ruptures, which correlates with lamin levels at the nuclear rim. The presence of persistent LEM-2 accumulation, and Nup160 loss, at rupture sites despite multiple cycles of nuclear entry and exclusion of GFP::α-tubulin suggest that lamin-dependent mechanisms exist that temporarily block rupture sites prior to full restoration of the NE.

### Dynein-dependent forces exacerbate nuclear envelope ruptures caused by lamin mutation

Next, we analyzed the role cytoskeletal forces play in lamin-dependent NE rupture and recovery. Prior work in cancer cells showed that actin-, and not microtubule-, derived forces induce ruptures at NE sites devoid of lamins (Hatch and Hetzer, 2016; Lammerding and Wolf, 2016). In *C. elegans* embryos, actin filaments are enriched on the anterior cortex of the embryo, away from the paternal pronucleus, at time points corresponding to NE ruptures (Figure S4A, Davies et al. 2015), suggesting that they likely do not directly impose forces on pronuclei to cause ruptures (Munro et al. 2004; Cowan & Hyman 2004; Oegema & Hyman 2006). On the other hand, the microtubule motor dynein is directly linked to the NE through the outer nuclear membrane LINC-complex protein, ZYG-12, and functions in separating NE associated centrosomes and positioning the paternal pronucleus at time points that correspond to NE rupture (Figure 1A, 2A, 2B, and Figure 3A, 3D (Gönczy *et al*., 1999; Malone *et al*., 2003; Oegema and Hyman, 2006; Luxton and Starr, 2014), suggesting that it may play a role in NE rupture or recovery. We eliminated centrosomes as the cause of NE ruptures because the maternal pronucleus, which is not directly associated with centrosomes (Cowan & Hyman 2004), also undergoes transient ruptures during this time period (Figure S4B, S4C). To test if dynein and NE-associated dynein are involved in NE rupture and recovery, we inhibited dynein and *zyg-12* and scored the percentage of pronuclear membranes that contained a Nup160::GFP gap. We found that percentage of pronuclei in lamin-inhibited embryos with Nup160::GFP gaps were unaffected by dynein or *zyg-12* RNAi-depletion (Figure 5A, 5B, Movies S4 and S5). Furthermore, the majority of dynein-inhibited and ZYG-12 inhibited lamin-N209K pronuclei that contained a Nup160::GFP gap also contained a condensed focus of chromatin adjacent to the Nup160::GFP gap (Figure 5A). Neither dynein nor lamin inhibition affected the transient recruitment of LEM-2 to NE rupture sites, which recruits repair machinery to NE holes (Figure S4D). Taken together, these data show that NE protein dynamics at NE rupture sites is unaffected by dynein-inhibition in lamin RNAi-depleted and lamin-N209K mutant embryos, suggesting that dynein forces imposed on the NE do not induce transient NE ruptures.

**Figure 5.**
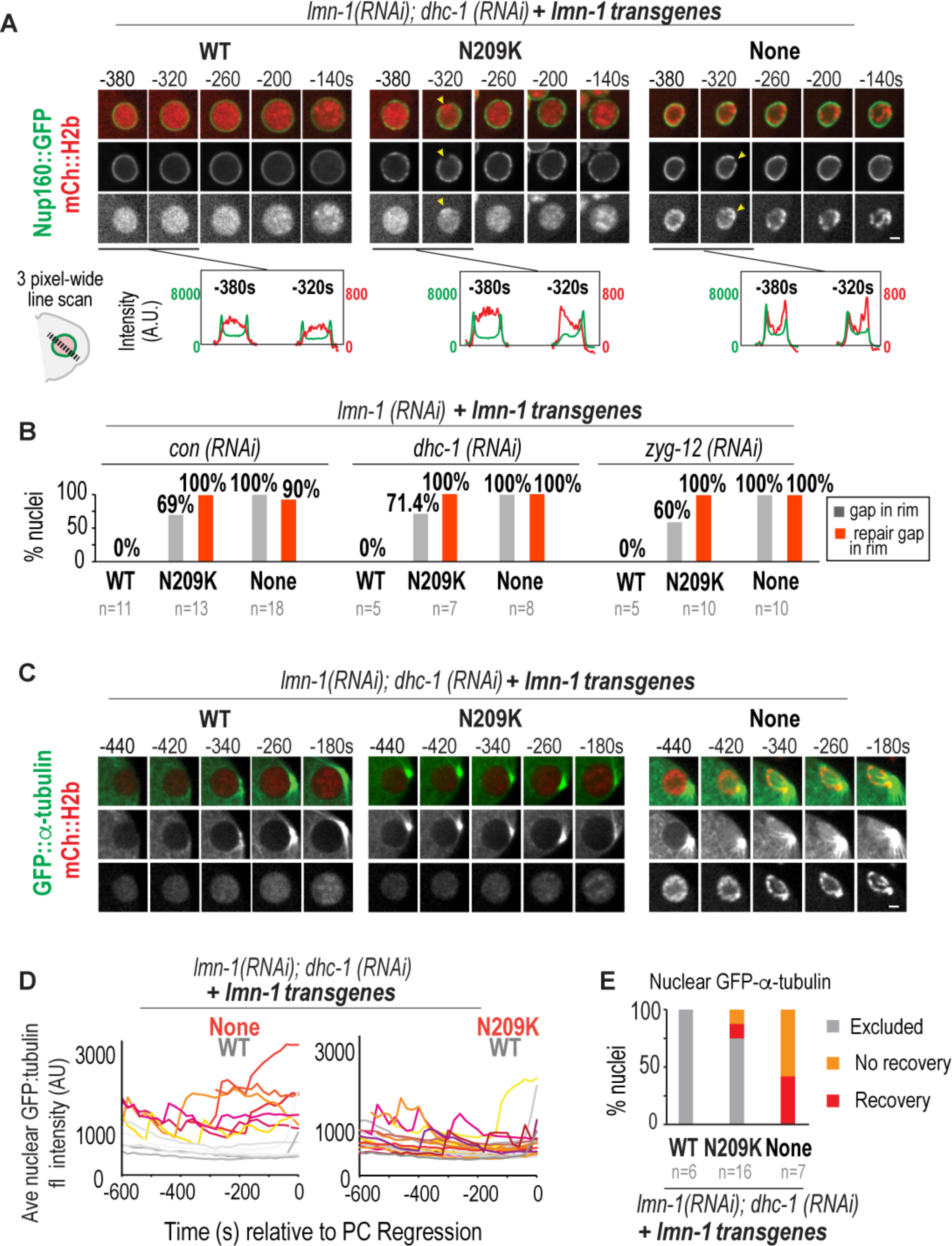
Dynein does not induce ruptures but exacerbates nuclear-cytoplasmic mixing in the presence of lamin-N209K. (A) (*Top*) Select images of paternal pronuclei expressing Nup160::GFP and mCherry::Histone2B from time-lapse series of lamin and dynein *(dhc-1)* co-depleted embryos in indicated lamin transgenic strains. Times are relative to PC Regression. Scale bar, 2.5μm. *(Below)* Graphs plotting intensities of Nup160::GFP (green) and mCherry::Histone2B (red) along 3 pixel wide line scan drawn across nucleus before and after gap formation for indicated conditions above. (B) Percent of paternal pronuclei with a Nup160::GFP gap (grey) and resolution (red) in the indicated conditions. n = number of embryos. (C) Select images of paternal pronuclei labeled with GFP::α-tubulin, and mCherry::Histone2B from time-lapse series of lamin and dynein co-depleted embryos in indicated lamin transgenic strains. Times are relative to PC Regression. Scale bar, 2.5μm. (D) Graphs plotting the average GFP::α-tubulin intensities over time in seconds relative to PC Regression for individual nuclei from indicated conditions. (E) Percent of nuclei from (D) that exclude GFP::α-tubulin (grey) or contain nuclear GFP::α-tubulin (yellow and red). Nuclei containing GFP:α-tubulin undergo recovery to <50% of maximum signal (red) or do not undergo recovery (orange) prior to pseudocleavage regression (−600s-0s). n = number of embryos. See also Figure S4, Movie S4, and Movie S5.

Despite the abnormal initial appearance of condensed chromatin foci adjacent to NE rupture sites, we noticed that overall chromatin organization was partially rescued in lamin-N209K embryos by dynein- and *zyg-12*-inhibition (Figure 5A, 5C, S4D, Figure 6A, Movie S4). Coincident with recovery of the Nup160::GFP gap at the nuclear rim, condensed chromatin focus at rupture sites dispersed and chromosome condensation advanced with similar kinetics and distribution as control embryos (Figure 5A, Figure 6A, Movie S4). In lamin RNAi-depleted embryos, defects in chromatin condensation and organization were unaffected by dynein- or *zyg-12*-inhibition, indicating that lamin’s function in regulating chromatin organization does not depend on dynein (Figure 5A, 5C, S4D, Figure 6A, Movie S5). Thus, while NE and chromatin dynamics at rupture sites are unaffected by dynein inhibition in lamin-N209K and lamin mutant nuclei, nuclear dynein inhibition rescues chromatin condensation defects induced by lamin-N209K. Based on these data, we suggest that chromatin condensation defects in lamin-N209K pronuclei are a consequence of NE rupture, and that dynein forces imposed on the NE exacerbate the extent of NE ruptures when lamins are weakened.

**Figure 6.**
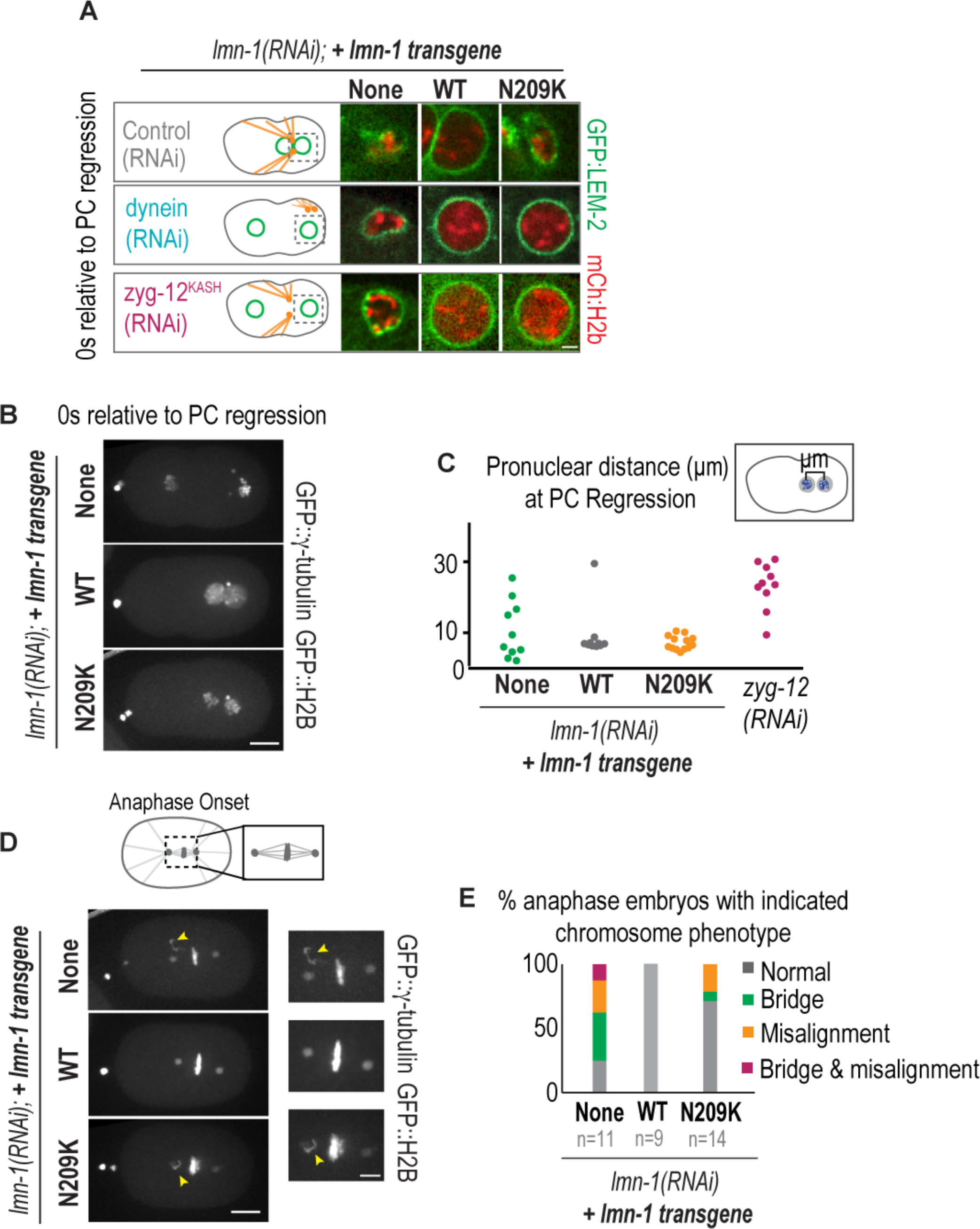
The lamin-N209 residue resists dynein forces during pronuclear migration to prevent loss of nuclear integrity and chromosome scattering prior to metaphase. (A) Confocal images of GFP::LEM-2 and mCherry::histone2B labeled paternal pronuclei at the time of pseudocleavage regression from indicated RNAi conditions in embryos expressing indicated transgenes. Scale, 2.5μm. (B) Maximum projection of confocal image stacks of GFP::γ-tubulin and GFP::Histone2B from lamin depleted *C. elegans* embryos expressing indicated transgenes. Scale bar, 10μm. (C) Graph plotting the distance in μms between the maternal and paternal histone masses as seen in (B) at PC Regression for the indicated conditions. (D) Maximum projection of confocal image stacks of GFP::γ-tubulin and GFP::Histone2B at anaphase onset from lamin depleted *C. elegans* embryos expressing indicated transgenes. Scale bar, 10μm. Insets are magnified images of metaphase plate at anaphase onset. Arrowheads indicate misaligned chromosome(s). Scale bar, 5μm. (E) Graph plotting the percentage of embryos with normal chromosome organization (grey), chromosome bridges (green), chromosome misalignment (yellow), or both (magenta) for the indicated conditions. n = number of embryos scored. See also Figure S5.

A prediction from the idea that lamin resists dynein forces to limit the extent of NE ruptures is that inhibition of dynein forces will reduce the accessibility of GFP::α-tubulin to the nuclear interior when lamin is present. In lamin RNAi-depleted embryos, GFP::α-tubulin accessed the nuclear interior even in the absence of dynein (Figure 5C-E). In contrast, dynein inhibition in lamin-N209K mutant embryos significantly reduced the percentage of embryos in which GFP::α-tubulin entered the nuclear interior (Figure 5C-E). Thus, dynein-induced tension on ruptured nuclear membranes interferes with lamin’s ability to prevent nuclear entry of GFP::α-tubulin. Depletion of lamin causes defects in the barrier function of the NE that are independent of the presence dynein forces.

We conclude that dynein-dependent tension on ruptured nuclear membranes enhances the severity of NE ruptures as indicated by the extent of defects in chromatin organization and nuclear access of GFP::α-tubulin in lamin-N209K mutant pronuclei inhibited of dynein. We propose that dynein forces counteract lamin-dependent mechanisms that function to restrict nuclear access of cytoplasmic proteins upon NE rupture.

### An essential role for lamin-N209 is to resist dynein forces during pronuclear migration to prevent chromosome scattering prior to metaphase

We predicted that the absence of pronuclear migration due to inhibition of dynein would allow us to track NEs with longer recovery periods in embryos with lamin perturbations. We first determined if dynein- or *zyg-12*-depletion rescues nuclear deformation and collapse upon pronuclear migration in lamin RNAi-depleted and lamin-N209K embryos. In lamin-N209K embryos, RNAi-depletion of dynein or *zyg-12* inhibits pronuclear migration and meeting and rescues pronuclear collapse, which occurs at the onset of pronuclear migration (Figure 6A). To confirm that rescue of pronuclear collapse was not because of a lack of penetrant depletion of endogenous lamin, we RNAi-depleted *zyg-12* or dynein in lamin-N209K mutant in the *lmn-1Δ* genetic background (Figure S5). Under these conditions, inhibition of dynein or *zyg-12* rescued early pronuclear collapse in lamin-N209K, and lamin-N209K nuclei broke down closer to the time of NEBD in WT-lamin embryos (Figure S5, see Figure 1D), indicating that dynein-dependent forces are the cause of pronuclear collapse in lamin-N209K embryos. In contrast, inhibition of dynein did not rescue nuclear deformation in lamin RNAi-depleted embryos and these nuclei retained nuclear GFP::α-tubulin (Figure 6A, Figure 5C-E). Thus, lamin is required to maintain NE integrity, even in the absence of dynein forces, and thus must be present for the NE to recover from damage. The N209 residue in lamin resists dynein forces to prevent irreversible nuclear collapse and is thereby essential to NE recovery from ruptures.

We noticed that despite pronuclear deformation and collapse during pronuclear migration in lamin-N209K embryos, the majority of chromosomes from each pronucleus maintain close proximity upon pronuclear meeting (Figure 6B, 6C). In 21% of cases, a few chromosomes scatter, leading to chromosome missegregation (Figure 6D, 6E). The distance between pronuclear masses is more severely affected in lamin RNAi-depleted embryos, which may be an outcome of defects in nuclear-centrosome attachment (Figure 6B, 6C; Meyerzon et al., 2009) and premature loss of NE integrity (Liu et al., 2000). Lamin RNAi-depleted embryos, but not lamin-N209K embryos, display a high percentage of chromosome-bridges (Figure 6E), a defect that has been described in prior work (Liu et al., 2000). These data suggest that while lamin is required for many processes in the early embryo, as has been suggested by others (Liu *et al*., 2000; Meyerzon *et al*., 2009; Hayashi, Kimura and Kimura, 2012), the main defect in the lamin-N209K mutant is to structurally protect NEs against dynein-dependent forces that enhance the severity of nuclear ruptures and induce nuclear collapse, which has downstream consequences on chromosome congression prior to anaphase.

We conclude that lamin is required to maintain nuclear integrity even in the absence of dynein forces that position pronuclei. In lamin-N209K embryos, nuclear collapse during pronuclear migration uncouples chromosome de-compartmentalization from spindle assembly leading to mitotic errors. The percentage of embryonic lethality (~35%) in lamin-N209K mutants depleted of endogenous lamin is similar to the percentage of chromosome bridges and missegregration defects in one-cell stage embryos (~28%, Figure S1B, 6E). In contrast, we find NE rupture sites form in ~70% in lamin-N209K embryos (Figure 3C), suggesting embryos survive to hatching despite transient losses of nuclear compartmentalization during early embryogenesis. We suggest that lethal defects in embryogenesis caused by expression of lamin-N209K are specific to the first division of the embryo in which pronuclei must migrate across the length of the embryo to mix their haploid genomes.

## DISCUSSION

Nuclear envelope rupture, which is distinct from nuclear collapse, occurs when a breach appears in the NE without nuclear membranes collapsing onto chromatin (Hatch & Hetzer 2014), and can be restored via ESCRT-mediated repair mechanisms (Denais *et al*., 2016; Raab *et al*., 2016). We provide the first mechanistic study of NE rupture and repair *in vivo*. Several observations support the claim that NEs rupture and repair in this system: 1) soluble GFP::α-tubulin, which is normally excluded from the nucleus, transiently enters the nucleus, 2) a single Nup160::GFP gap forms and resolves at the nuclear rim, 3) the NE transiently deforms upon Nup160::GFP gap formation, 4) the integral NE protein LEM-2 accumulates and subsequently disperses in a confined region at the nuclear rim, and 5) locally condensed chromatin protrudes precisely at the NE site with accumulated LEM-2 foci and that is devoid of Nup160::GFP (Figure 2-6). By analyzing a partial-loss-of function lamin mutation in the first cell division of the transcriptionally quiescent *C. elegans* zygote, we demonstrate that by counteracting dynein induced tension on the NE lamin promotes a distinct step in NE recovery that involves restricting nuclear access to cytoplasmic proteins prior to full restoration of the NE (Figure 7). Further, we provide evidence that mechanisms of recovery from NE rupture exist in wildtype *C. elegans* embryos since NEs recover from loss of the permeability barrier caused by laser-induced puncture of the NE. Taken together, the relatively short timescale of transient NE ruptures (~100-200s) in the *C. elegans* zygote allowed us to define the dynamics of NE reorganization at rupture sites during NE rupture and recovery, to correlate these events to cell cycle progression, and to recapitulate the process in wild type embryos. In this section, we discuss the opposing roles for lamin and dynein during NE rupture and recovery and compare the dynamics of NE reorganization at rupture sites to mechanisms of NE reformation at the end of mitosis.

**Figure 7.**
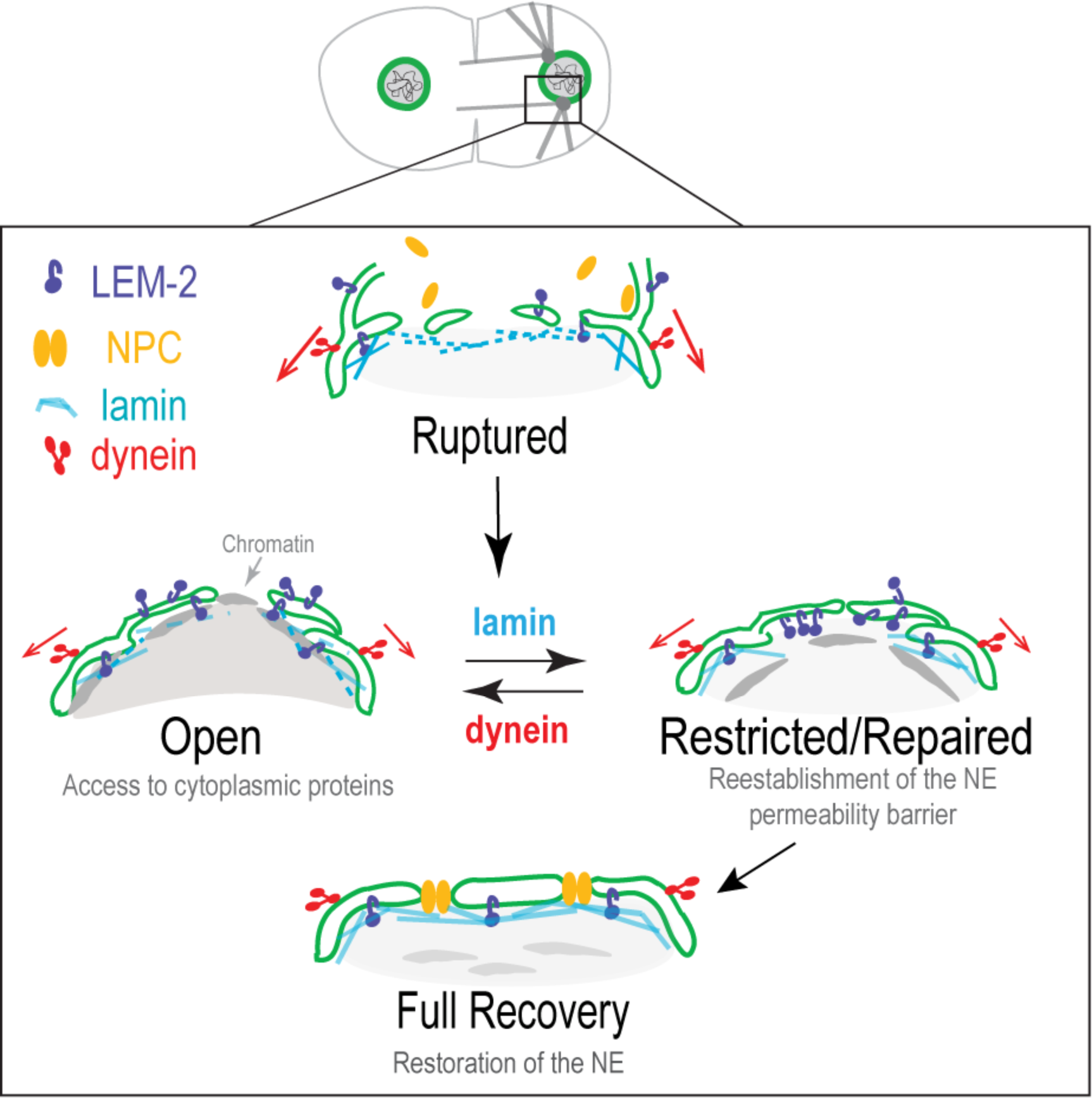
Lamin resists dynein forces on ruptured nuclei to restrict nuclear access to cytoplasmic proteins and promote NE recovery. Lamin (cyan) inhibition induces NE ruptures leading to clearance of NPCs (yellow) from NE rupture sites. Dynein forces (red) impose tension on ruptured nuclear membranes, which exacerbates nuclear cytoplasmic mixing by enabling access to cytoplasmic proteins (“Open”). Lamin resists dynein forces to promote a “Restricted/Repaired” state that prevents further access to cytoplasmic proteins and allows reestablishment of the permeability barrier. “Full Recovery” of the NE is a distinct step that leads to restoration of the NE through reinsertion of NPCs to the healed region of the NE, dispersal of LEM-2 from rupture sites, and reestablishment of chromatin organization.

### Lamin resists dynein forces to restrict loss of the nuclear permeability barrier following nuclear envelope rupture

We propose a model in which lamin resists dynein forces imposed on the NE to modulate the extent of loss of the nuclear permeability barrier caused by rupture and the ability of ruptured nuclei to recover (Figure 7). In our model, dynein-generated forces pull ruptured nuclear membranes apart, while lamin assembly opposes these forces to restrict access to cytoplasmic proteins by keeping membranes in close proximity and preventing repeated ruptures at these sites. Lamin depletion induces NE ruptures in the absence of dynein forces, suggesting that dynein forces are not required to cause ruptures. Instead, we suggest that dynein-generated tension on the NE impacts the severity of NE ruptures. In support of this idea, we show that dynein-inhibition rescues detectable nuclear entry of GFP::α-tubulin in lamin-N209K mutant nuclei, yet our data suggest that these nuclei still undergo ruptures as indicated by the unchanged frequency of Nup160::GFP gaps and the transient presence of condensed chromatin adjacent to these gaps. The transient accumulation of GFP::LEM-2 foci at the nuclear rim suggests that these regions recruit repair machinery (Gu et al. 2017; Webster et al. 2016), which provides further evidence that NE holes exist at these sites. This idea is supported by work in migrating cancer cells showing that recruitment of repair machinery to NE rupture sites is unaffected by lamin inhibition (Celine M Denais et al. 2016), whereas the duration of NE ruptures is lamin-dependent (Vargas et al. 2012; Denais et al. 2016; Raab et al. 2016). Because the localization and dynamics of Nup160 at NE ruptures sites is unaffected by dynein inhibition, we suggest that Nup160 organization at rupture sites is regulated by a distinct mechanism than that which has been described for dynein and lamin in NPC organization (Guo & Zheng 2015). Thus, our time-lapse data of NE reorganization and protein dynamics in lamin inhibited nuclei provides evidence for a lamin-dependent mechanism that restricts cytoplasmic-nuclear mixing upon rupture (Figure 7). We suggest that lamin’s role in restricting loss of the nuclear permeability barrier upon rupture is to counteract cytoskeletal forces that continue to generate forces on the weakened NE thereby preventing repeated breaks at weakened rupture sites.

We were unable to monitor lamin localization at NE rupture sites in our system, however work in cell culture shows that NE rupture sites are devoid of lamin B1 (De Vos et al. 2011; Vargas et al. 2012; Maciejowski et al. 2015; Denais et al. 2016), and that LINC-complex components do not localize to NE regions devoid of lamins (Hatch and Hetzer, 2016), suggesting that these sites are not subject to direct tension generated by the cytoskeleton. This work showed that regions devoid of lamin B1 are present even in the absence of nuclear-cytoplasmic mixing, which supports our hypotheses that nuclear ruptures may be temporarily occluded prior to repair or that these sites are repaired but prone to reoccurring ruptures. One possibility is that upon NE rupture, unknown mechanisms, which we hypothesize are ER-mediated, form a temporary barrier and that cytoskeletal forces that generate tension on the nucleus outside of this site disrupt the formation or maintenance of this barrier, which is devoid of lamins and therefore susceptible to damage (Figure 7). It is also possible that lamin assembly at rupture sites follows repair causing these sites to be more likely to rupture again. Our model suggests that the remaining lamin meshwork at intact regions of the nuclear periphery maintain NE integrity following NE rupture by resisting cytoskeletal forces (Figure 7). During pronuclear positioning in the *C. elegans* zygote, lamin resists forces generated by dynein motors that continue to directly pull on intact regions of the NE (Gönczy et al. 1999; Kimura & Onami 2005; Malone et al. 2003; Oegema & Hyman 2006); these forces tear apart rupture sites and in turn increase the severity and duration of ruptures. Thus, we identified a role for lamin in resisting dynein forces to limit the severity of NE ruptures that is likely important across different cell types that utilize dynein-dependent mechanisms to position nuclei (Coffinier et al. 2011; Zhang et al. 2009; Bone & Starr 2016). We suggest that any source of tension on the NE upon rupture requires lamin assembly at sites outside of ruptures to prevent increased stress on NE holes and in turn restrict the severity of nuclear-cytoplasmic mixing prior to repair.

### Mechanisms that induce nuclear envelope rupture in the early *C. elegans* embryo

In cells in culture, constriction during cell migration or actin-derived compression forces cause nuclear herniations in regions devoid of lamins (Harada et al. 2014; Denais et al. 2016; Hatch & Hetzer 2016; Raab et al. 2016; Thiam et al. 2016; Irianto et al. 2017), which increases the surface tension. To release this tension, the NE ruptures at these sites (Denais *et al*., 2016; Raab *et al*., 2016; Thiam *et al*., 2016b; Irianto *et al*., 2017). Furthermore, not all rupture sites correlate with chromatin herniations, suggesting that herniations are not a prerequisite for rupture (Hatch & Hetzer 2016). We did not detect chromatin herniations upon NE rupture in *C. elegans* pronuclei. Instead, we found that condensed chromatin puncta appear at rupture sites – an observation also made in lamin A deficient mammalian cells (Robijns et al. 2016) and in migration-induced ruptures (Irianto et al. 2016) - several time points after rupture, which correlated with entry into mitosis. This delayed appearance of condensed chromatin relative to the appearance of other indicators of rupture suggests that chromatin puncta are a consequence rather than a cause of rupture.

Our data show that dynein, which generates tension on the order of tens of piconewtons on pronuclei at time points that correspond to rupture (Kimura & Onami 2005), is not required to induce NE ruptures. Actomyosin contractile forces localize at the cortex far from centrally located nuclei and thus are not directly coupled to the NE (Cowan & Hyman 2004; Munro et al. 2004; Oegema & Hyman 2006). However, we cannot exclude the possibility that direct or indirect actin-generated forces, such as cytoplasmic flow generated by asymmetric actomyosin contractility (Hird & White 1993), induce NE ruptures in this system. Interestingly, micronuclei with reduced lamin undergo ruptures independent of compression forces, supporting the idea that in specific contexts disorganization of the nuclear lamina is sufficient to induce NE ruptures independent of cytoskeletal forces (Hatch et al. 2013). Further studies are required to determine if lamin inhibition on its own or in combination with internal or external pressure imposed on the nucleus induces ruptures.

### Local chromatin and nuclear envelope reorganization during nuclear envelope rupture and recovery

Our works in *C. elegans* shows that the region devoid of Nup160::GFP is ~800nm to 3μm in the xy directionwhen the paternal pronucleus is ~4μm to 7μm in diameter, which we suggest is likely significantly larger than the size of NE holes. This idea is supported by the relatively longer recovery times following laser-induced punctures of the nuclear rim that had dimensions of ~1μm to 2μm in the y-direction and ~0.5μm in the z. In addition to clearance of NPCs from rupture sites, the inner nuclear membrane protein LEM-2 accumulates in distinct regions on the nuclear rim at the time of rupture. Recent work showing a conserved role for Lem2p/LEM2 in yeast and mammalian cells in recruiting components of the ESCRT machinery to NE holes suggests that LEM-2 foci mark sites of NE repair (Gu et al. 2017; Webster et al. 2016). The removal of NPCs and accumulation of LEM-2 at sites of NE rupture resemble mechanisms related to NEBD and reformation. An intriguing future direction is to understand the molecular trigger that induces the downstream reorganization of the NE to promote NE reformation upon NE rupture.

In addition to the reorganization of NE proteins upon rupture, it is compelling that compacted chromatin foci form directly adjacent to rupture sites upon entry into prophase. We suggest that chromatin may be more readily accessible to mitotic-activated cytoplasmic factors, which results in its accelerated condensation relative to distally located nuclear chromatin. Alternatively, local reorganization of NE proteins induced by rupture may alter the distribution of condensed chromatin. Interestingly, penetrant loss of lamin results in peripherally associated chromatin throughout the nucleus. Depletion of nucleoporins Nup205/93, which are required to set the exclusion limit of the nuclear permeability barrier, also causes association of chromatin to the nuclear periphery in *C. elegans* embryos (Galy et al. 2003). Thus, impaired NPC function in lamin depleted pronuclei suggests that although rupture sites recover, the permeability barrier function of the NE that depends on lamin does not. Local loss of lamin and NPCs at rupture sites in lamin-N209K nuclei may cause the transient association of chromatin to this site via an unknown mechanism that involves reorganization of NE associated proteins.

### The relationship between transient nuclear ruptures and the cell cycle

It has been suggested that NE rupture and repair mechanisms are restricted to interphase (Hatch & Hetzer 2014). We found that repair mechanisms active during early prophase in the *C. elegans* zygote ensure maintenance of NE integrity during pronuclear positioning. Tissue culture cells couple mitotic-entry to NEBD, whereas the *C. elegans* zygote undergoes an elongated prophase (~450s) upon entry into mitosis, which coincides with dynein-dependent pronuclear positioning (Oegema & Hyman 2006). Nuclear envelope repair mechanisms may be inactivated by factors that trigger NEBD-mediated NE permeabilization, which coincides with diffusion of INM proteins into the ER. One possibility is that diffusion of LEM-2 into the ER upon NE permeabilization provides a mechanism to inhibit NE repair during this time by removing the nuclear specific adaptor for the ESCRT complex. Subsequent to NE permeabilization, disassembly of phosphorylated nuclear lamins permits dynein-dependent forces to remove permeabilized nuclear membranes from chromatin (Salina et al. 2002), which is analogous to the opposing roles we identified for lamin and dynein on transiently ruptured nuclear membranes. We speculate that other mechanistic similarities exist between this process and mitotic regulated NEBD and reformation and that a key distinction will be in the signaling pathways that restrict nuclear-cytoplasmic mixing upon rupture to prevent downstream consequences in genomic integrity prior to mitotic-coupled NEBD.

## ACKNOWLEDGEMENTS

We thank Patrick Lusk, Megan King, Topher Carroll, Joshua Gendron, and Julie Canman for insightful discussions, experimental suggestions, and helpful critiques. Arshad Desai, Karen Oegema and Julie Canman for strains used in experiments. We thank our sources of support from Yale University and the Ludwig Institute for Cancer Research (Ronald Biggs). The National Institute of Health for support of Lauren Penfield, (T32 GM007499) and Michael Mauro (T32GM007223-S1). Daniel Needleman was supported in part by the National Institute of Health (1R21HD080057-01A1) and the National Science Foundation (DMR-0820484).

